# From group to individual - Genotyping by pool sequencing eusocial colonies

**DOI:** 10.1101/2021.11.08.467442

**Authors:** Sonia E Eynard, Alain Vignal, Benjamin Basso, Yves Le Conte, Axel Decourtye, Lucie Genestout, Emmanuelle Labarthe, Fanny Mondet, Kamila Tabet, Bertrand Servin

## Abstract

**Background:** Eusocial insects play a central role in many ecosystems, and particularly the important pollinator honeybee (*Apis mellifera*). One approach to facilitate their study in molecular genetics, is to consider whole colonies as single individuals by combining DNA of multiple individuals in a single pool sequencing experiment. Such a technique comes with the drawback of producing data requiring dedicated analytical methods to be fully exploited. Despite this limitation, pool sequencing data has been shown to be informative and cost-effective when working on random mating populations. Here, we present new statistical methods for exploiting pool sequencing data of eusocial colonies in order to reconstruct the genotype of the colony founder, the queen. This leverages the possibility to monitor genetic diversity, perform genomic-based studies or implement selective breeding.

**Results:** Using simulations and honeybee real data, we show that the methods allow for a fast and accurate estimation of the genetic ancestry, with correlations of 0.9 with that obtained from individual genotyping, and for an accurate reconstruction of the queen genotype, with 2% genotyping error. We further validate the inference using experimental data on colonies with both pool sequencing and individual genotyping of drones.

**Conclusion:** In this study we present statistical models to accurately estimate the genetic ancestry and reconstruct the genotype of the queen from pool sequencing data from workers of an eusocial colony. Such information allows to exploit pool sequencing for traditional population genetics, association studies and selective breeding. While validated in *Apis mellifera*, these methods are applicable to other eusocial hymenoptera species.

## Introduction

Eusocial organisms such as bees, ants or wasps live in large colonies produced by a single individual (the queen) and have a specific mating system in which the queen is mated to a cohort of males. In the case of the honeybee, *Apis mellifera*, a colony is typically composed of a single queen, a large number (up to tenths of thousands) of workers and a few males. The queen is usually the only reproducing individual and all individuals present in the colony are its offspring. In the wild, after mating with a cohort of 10 to 20 males the virgin queen will return to the colony and maintain its population, throughout her life, by continuously laying eggs. Fertilised eggs will produce diploid worker females, while unfertilised eggs will produce haploid males. Males are therefore a direct sample of the queen genome and can be considered as flying gametes. The mosaic composition of a colony makes standard genomics analysis complex especially when making breeding decisions (Brascamp and Bijma, 2014; Uzunov, Brascamp, and Büchler, 2017). In eusocial populations, each worker performs individual tasks participating in the collective phenotype of the colony. However, although the phenotype of the colony is collective, the queen contributes to more than half of the genetics of the colony (through diploid female and haploid male offspring) that will be passed on to next generations. Thus, the queen’s genotype itself is an essential piece of information for genetic analysis aimed at studying the evolution of populations or performing selective breeding. Even though the field of insect genomics has boomed in the past decades there still is a need to expand traditional approaches of population genetics for this specific kind of organisms (Toth and Zayed, 2021). However, contrary to large animal species, sampling the queen for genotyping is impossible without threatening its integrity and is therefore rarely performed in routine beekeeping practices. One possible approach to overcome these problems is to perform individual or pool genotyping (Petersen et al., 2020) of a set of males. However this implies an increased manipulation effort to sample the individual males or sequencing cost as multiple genotyping experiments are required to infer the genotype of a single queen.

Advances in sequencing technologies have brought new opportunities to develop tools for genomics and genetics. Amongst these, parallel sequencing allows for counting of sequencing reads at all positions on the genome which thus permitted the development of pool sequencing for allele frequencies estimations (Schlotterer, Tobler, Kofler, and Nolte, 2014). By combining DNA from multiple individuals into a unique sequencing experiment, pool sequencing allows for cheap and fast data acquisition, especially for non-model organisms for which resources are limited. However pool sequencing outcomes, allele counts in the pool instead of genotypes, are more difficult to use in practice and require specific programs and software to perform SNP calling, mainstream population genetics analysis, association testing (Kofler, Pandey, and Schlötterer, 2011; Bansal, 2010; Purcell et al., 2007; Chang et al., 2015; Zhou and Stephens, 2012; Speed, Holmes, and Balding, 2020) and more. Additionally traditional pool sequencing is performed on a group of unrelated individuals representing a population often linked by an environmental factor (e.g a population in a specific location, a genetic type …).

In this study, we propose a new application of pool sequencing to multiple individuals from a single colony in the context of eusocial insects. Hence, contrastingly to standard pool experiment, representing a population of individuals, pool experiment on colonies can be seen as sequencing of a meta-individual. Using this specificity we introduce dedicated statistical methods to estimate the genetic ancestry of the queen and reconstruct its genotype from pool sequencing of workers. The acquisition of genotype data will on the one hand provide information on the queen that can further benefit breeding decisions and will on the other hand allow the use of standard programs and software for population genetics analysis such as admixture or association studies. Two models are proposed and evaluated: the first model estimates the genetic ancestry of the queen, based on single colony data but assuming information on the allele frequencies of markers in reference populations and the second model exploits information available across multiple colonies to reconstruct the queen genotype. Performances of the models are evaluated through simulations including some based on real data from a diversity panel in *Apis mellifera* (Wragg et al., 2021). Using these simulations we show that the genetic ancestry of the queen estimated from the pool sequencing data matches results from standard population genetics methods results on genotype data and that the genotype of the queen can be reconstructed with an error rate limited to a few percent. To evaluate the interest of pool sequencing compared to individual genotyping, we applied our genotype reconstruction models to real data in this species from a field experiment where both pool sequences of workers and individual sequences of male offspring from the same colony were available. We showed that inference of the genetic ancestry and the genotype of the queen based on pool sequencing data matches results obtained from individual data on male offspring.

Models introduced in this study can be used sequentially to first estimate the genetic ancestries of a population of colonies, then use this information to cluster the dataset into homogeneous populations and finally infer genotypes of colonies by considering them jointly within these homogeneous clusters. Finally we discuss the interpretation of the results obtained with the models proposed, their applicability and possible extensions.

## Materials and Methods

For the sake of understanding statistical models are presented here from the most simple to the most complex even though they can be used independly in the rest of the paper.

### Models

We consider data coming from colony pool sequencing experiments. For each colony, whole genome sequencing is assumed to be performed on DNA extracted from the mix of a large number of worker bees. For a colony *c*, the raw data consist of the reference allele counts and sequencing depths at a fixed set of *L* biallelic loci. At a locus *l*, with observed reference allele count 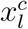 and sequencing depth 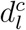, we have:

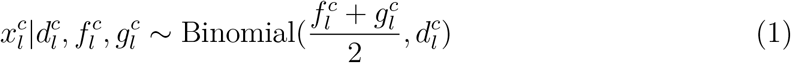

where 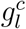 is the (unknown) queen genotype expressed as the frequency of the reference allele (*i.e*. 0, 0.5 or 1) and 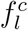 is the (unknown) reference allele frequency in the males that mated with the queen. We are interested in reconstructing information on the possible genotypes of the queen 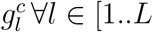. As 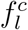 and 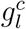 both contribute to the allele counts in the pool, it is clear that these parameters are unidentifiable without more information. To separate them, we thus need external information on 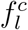 and/or 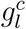. We now discuss two possibilities to incorporate such information and the associated inferences that can be drawn.

#### Homogeneous Population Model

In this approach, we add to model (1) the hypothesis that queens and males of all colonies come from the same random mating population. Under this hypothesis, (i) the allele frequency at a given locus is the same for all colonies and (ii) genotypes at a locus are sampled according to this frequency, so we have for a locus *l* :

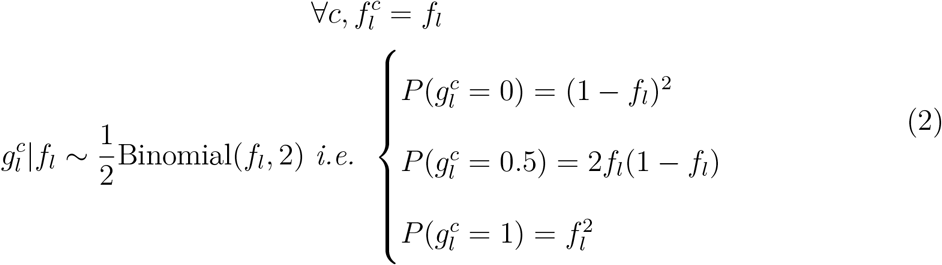

This new model has only one parameter per locus (*f_l_*) and the likelihood is:

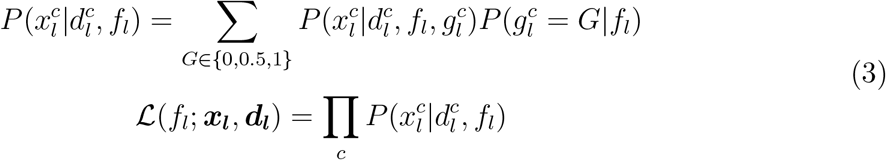

where ***x_l_*** is the vector of reference allele counts in all colonies and ***d_l_*** the corresponding vector of sequencing depths. The likelihood (3) is maximized numerically for *f_l_* on [0, 1]. The maximizing value (called the Maximum Likelihood Estimate, MLE) 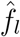 can be used for inference on ***g_l_*** based on the posterior distribution 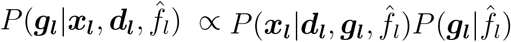.

This homogeneous population model (HP) should only be applied when the set of colonies have a similar genetic background. We therefore developed another approach, the admixture model, aimed at estimating the genetic ancestry of a single colony from pool sequencing data.

#### Admixture Model

The objective of this model is to describe the “genetic background”, the subspecies, of a colony. To do so, we will adopt the widely used modeling framework introduced by Pritchard, Stephens, and Donnelly (2000) and define the genetic background of a colony as *the proportions of the queen genome that come from a set of pre-defined reference populations* (in our applications below, the reference populations considered are *Apis mellifera mellifera, Apis mellifera ligustica & carnica* and *Apis mellifera causasia*, the three main populations found in Western Europe (Wragg et al., 2021)). We will do that in a supervised manner so we will assume that we are provided with allele frequencies in a set of *K* reference populations at the *L* loci : this takes the form of an *L* × *K* matrix ***F*** where *F_lk_* is the frequency of the reference allele at locus *l* in population *k*. Here we are interested in inferring ***q***, the *K*-vector of admixture proportions for the queen: *q_k_* is the proportion of alleles over all loci that come from population *k*. Dropping the *c* index as the model is fitted for each colony independently, the likelihood for ***q*** is:

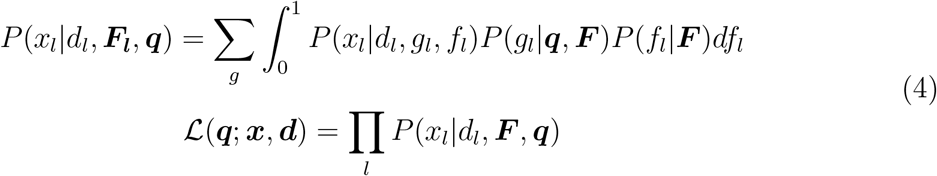

In order to compute likelihood (4), we need to specify *P*(*g_l_*|***q, F***), the prior distribution on *g_l_* given the admixture proportions, and *P*(*f_l_*|***F***) the prior on the allele frequency at locus *l*. To perform inference we need to devise a way of maximizing the likelihood (4). We now explain how we addressed these two issues.

#### Priors

To specify the prior *P*(*g_l_*|***q, F***), we use the classical approach of introducing latent variables 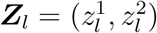 at each locus *l* that denotes the origins (in terms of reference populations) of the two alleles carried by the queen. Then we can write:

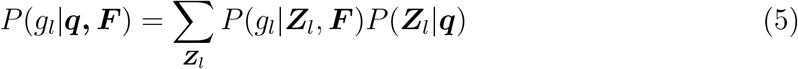

where *P*(*g_l_*|***Z_l_, F***) is the probability of the queen genotype given the origins of the two alleles, which is a function of the allele frequencies in the *K* reference populations (*e.g*. *P*(*g_l_* = 0.5|***Z_l_*** = (2, 2), ***F***) = 2*F*_2*l*_(1 – *F*_2*l*_)), and *P*(***Z_l_***|***q***) is the probability of the pair of origins that depends on the admixture proportions ***q*** (e.g. 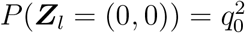).

For *P*(*f_l_*|***F***), the prior on the allele frequency in males mated to the queen, we use an informative prior based on the allele frequencies in the reference populations:

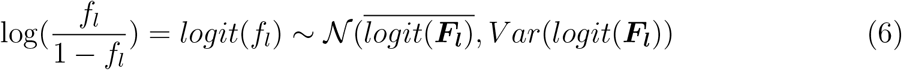

This prior is informative if all reference populations have similar allele frequencies and more diffuse if allele frequencies in reference populations differ greatly. Finally, the estimation of the vector ***q*** is performed using an EM algorithm. Note that this is similar to the supervised version of the estimation procedure of the Pritchard et al. (2000) model as the matrix of allele frequencies ***F*** is considered known a priori.

### Simulations

To evaluate the performance of the two models, we simulated data as obtained from a pool sequencing experiment. We assume these data come in the form of the reference allele counts 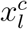 and sequencing depths 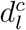 at each locus *l*, knowing the queen genotypes 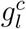 and allele frequencies in the inseminating drones 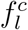. To further condition our simulations on what can be expected from real data, we exploited information available in a reference population of *Apis mellifera* (Wragg et al., 2021). This data consists of 628 European samples of haploid drones (Supplementary Table ST2) with genotypes available at 6,914,704 Single Nucleotide Polymorphisms (SNPs). Wragg et al. (2021) showed that this panel is structured into three main genetic background for which unadmixed (reference) individuals can be identified, with a threshold of 99% of their genetic background being from a unique type: the **M** background (*Apis mellifera mellifera*) with 85 reference individuals, the **L** background (*Apis mellifera ligustica & carnica*) with 44 reference individuals and the **C** background (*Apis mellifera caucasia*) with 16 reference individuals (Supplementary Table ST3). In the simulations described below, the reference panel information used was either the allele frequencies in the three main backgrounds (***F*** = (*F_lp_*) ∈ [0, 1]^*L*×3^, where the columns contain the allele frequencies of all *L* markers in genetic backgrounds L, M and C in this order) and/or the genotypes of the reference individuals.

#### Independent markers

To evaluate the performance of the models proposed, a first set of simulations was performed on 1000 independent SNPs chosen to be common and ancestry informative with respect to the L, M and C genetic backgrounds. To this goal, the 1000 SNPs were randomly sampled from the 722,170 SNPs out of the 6,914,704 that had a minor allele frequency (MAF) ≥ 0.1 and a variance across genetic backgrounds ≥ 0.1. For this first set of simulations, only the allele frequencies in the reference panel at the 1000 SNPs were used.

First, for each colony *c* the proportions of the genome coming from each of the genetic backgrounds (termed *genetic ancestry* from now on) of the queen 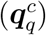 and the inseminating drones 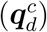 were sampled from a Dirichlet distribution:

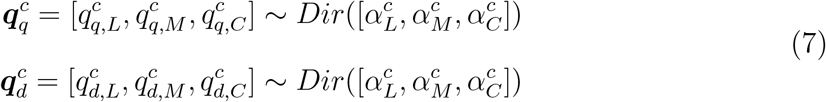

Different values were considered for the *α* parameters to consider different levels of admixed ancestries for the colony (Table 1). Simulated genetic ancestries are represented in Figure S1.

**Table 1:** Simulated genetic ancestries for queen and drones under Dirichlet distribution.

| Number of simulated colonies | Queen genetic ancestry | Dirichlet alpha parameters for queen | Drones genetic ancestry | Dirichlet alpha parameters for drones |
| --- | --- | --- | --- | --- |
| 100 | LMC | 10,10,10 | LMC | 10,10,10 |
| 40/30/30 | L___/___M___/___C | (10,0.5,0.5)/(0.5,10,0.5)/(0.5,0.5,10) | LMC | 10,10,10 |
| 40/30/30 | L___/___M___/___C | (10,0.5,0.5)/(0.5,10,0.5)/(0.5,0.5,10) | L___/___M___/___C | (10,0.5,0.5)/(0.5,10,0.5)/(0.5,0.5,10) |
| 100 | LM___ | 10,10,0.5 | LMC | 10,10,10 |
| 100 | LM___ | 10,10,0.5 | LM___ | 10,10,0.5 |
| 100 | L___ | 10,0.5,0.5 | L___ | 10,0.5,0.5 |
| 50/50 | L___/___M___ | (10,0.5,0.5)/(0.5,10,0.5) | L___/___M___ | (10,0.5,0.5)/(0.5,10,0.5) |
| 100 | ___MC | 0.5,10,10 | LMC | 10,10,10 |
| 100 | ___MC | 0.5,10,10 | ___MC | 0.5,10,10 |
| 100 | ___M___ | 0.5,10,0.5 | ___M___ | 0.5,10,0.5 |
| 50/50 | ___M___/___C | (0.5,10,0.5)/(0.5,0.5,10) | ___M___/___C | (0.5,10,0.5)/(0.5,0.5,10) |
| 100 | L___C | 10,0.5,10 | LMC | 10,10,10 |
| 100 | L___C | 10,0.5,10 | L___C | 10,0.5,10 |
| 100 | ___C | 0.5,0.5,10 | ___C | 0.5,0.5,10 |
| 50/50 | L___/___C | (10,0.5,0.5)/(0.5,0.5,10) | L___/___C | (10,0.5,0.5)/(0.5,0.5,10) |

Description of the simulations for colony and population size, composition and genetic ancestries. For each of the 15 scenarios designed for simulations we present the number of simulated colonies, the queen's genetic ancestry in term of genetic backgrounds *Apis m. caucasia* C, *Apis m. ligustica* & *carstica* L and *Apis m. mellifera* M, the associated Dirichlet alpha vectors, and the same information for the inseminating drones.

Second, the allele frequencies of each SNP *l* in the cohort of inseminating drones was simulated as:

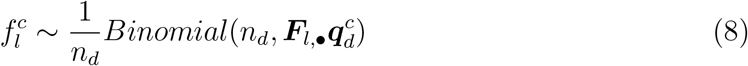

where ***F***_*l*,•_, is the *l*-th line of the ***F*** matrix and *n_d_* is the number of inseminating drones, here fixed at 15 (Tarpy and Nielsen, 2002; Tarpy, Nielsen, and Nielsen, 2004).

Third, the genotype of the queen at a SNP *l* was simulated by first drawing the population of origin of each of the two allele of the queen 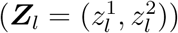 from a multinomial distribution with parameter 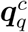. The genotype of the queen was finally obtained as 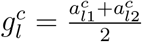 where :

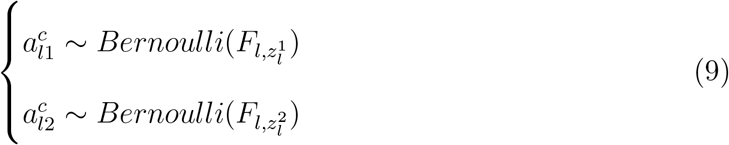

Finally, pool sequencing data was simulated as

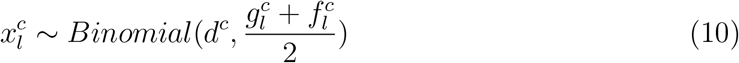

where *d_c_* is the sequencing depth, which was fixed at 30 unless otherwise specified in the Results section.

#### Linked markers

Pool sequencing experiments provide information on a large number of markers distributed throughout the genome. In order to evaluate the performance of the models in realistic conditions for the distribution of allele frequencies and the genetic structure, a second set of simulations was performed using individual genotypes of 628 individuals from the diversity panel previously described in Wragg et al. (2021) and used beforehand to define reference genetic backgrounds. First, individuals were clustered into seven groups, of all potential combinations of admixture between the three genetic backgrounds, using hard thresholds on their initial vectors of genetic ancestry estimated with ADMIXTURE (Alexander, Novembre, and Lange, 2009) (Figure S2). Then, each colony was simulated by sampling haploid genotypes of 17 individuals two of which were united to create the genotype of the queen (replacing step (9) above) and the remaining 15 were used as inseminating drones under different scenarios of admixture between the three populations, replacing step (8). Then pool sequencing data 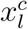 was simulated as in (10). The simulated scenarios are the same as for independent markers, despite that only 20 colonies are simulated per scenario because of sampling limitation due to the restricted number of individuals to select from. As an example, when the queen of the colony is L genetic background and the inseminating drones are LMC genetic background the two individuals to make the queen were sampled from the group of ’pure’ L and the 15 inseminating drones were sampled from all the possible groups, as their combinaison will create a mixture of genetic backgrounds.

##### Evaluation of statistical models

###### Genetic ancestry

For each colony and for each set of simulations, the queen genetic ancestry *q^c^* was estimated using the Admixture model (AM). For independent marker simulations, the estimates were compared to the true simulated value, while for linked marker simulations they were compared to the estimates obtained by running ADMIXTURE on the queen genotype. All simulated colonies were analysed jointly with AM and thereafter clustered into seven groups based on their ancestry vectors. Hence, each cluster was a group of colonies with homogeneous genetic ancestry.

###### Genotype reconstruction

The HP model was used to reconstruct the queen genotype, within each of the ancestry clusters described above, in the linked marker simulations. Criteria for evaluating the model were :

- the genotyping error rate measured as the proportion of errors in the reconstructed genotypes among all markers. We measured the genotyping error rate for different calling probability thresholds (see Results).
- the calibration of the posterior genotype probabilities. For each locus and each simulated colony, the HP model provides the posterior probabilities of the three possible genotypes. Because in the simulations the true genotype is known, we can evaluate in which proportion of the simulations (π) a genotype with posterior probability P is the true genotype. If the model is perfectly calibrated π = *P*. Hence, the calibration of the model was measured as

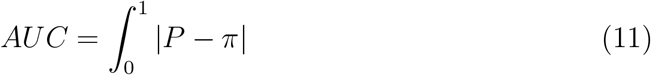 In practice we estimated π by grouping genotype probabilities in bins of size 0.05.

### Validation on experimental data

#### Dataset

In order to evaluate the performance of the genotyping by pool sequencing approach, we produced a new dataset where colonies were both pool sequenced and individual drones were sampled. Thirty four colonies, present at an experimental apiary and representing the diversity of French honeybee populations, were sampled in 2016. For each colony between approximately 300 and 500 worker bees were collected and pooled for sequencing purposes. DNA extraction was performed from a blended solution of all the workers of the colony with 4 m urea, 10 mm Tris-HCl pH 8, 300 mm NaCl, 10 mm EDTA. The elution was centrifuged for 15 min at 3500 g, and 200 μl of supernatant was preserved with 0.5 mg proteinase K and 15 μl of DTT 1 m for incubation overnight at 56 °C. After manual DNA extraction and DNA Mini Kit (Qiagen) a volume of 100 μl was used to perform pair-end sequencing on the IlluminaTM HiSeq 3000 or NovaSeq 6000 platform with the aim to obtain approximately 30× raw sequencing data per sample. Raw reads were then aligned to the honeybee reference genome Amel HAV3.1, Genebank assembly accession GCA_003254395.2 (Wallberg et al., 2019), using BWA-MEM (v0.7.15; (Li, 2013)). For pool sequenced experiments the resulting BAM files were converted into pileup files using Samtools mpileup (Li and Durbin, 2009) with the parameters: -C 50 coefficient of 50 for downgrading mapping quality for reads with excessive mismatches, -q 20 minimum mapping quality of 20 for an alignment, -Q 20 and minimum base quality of 20, following standard protocols. This procedure was applied exclusively to the 6,914,704 Single Nucleotide Polymorphisms (SNPs) identified in Wragg et al. (2021) as polymorphic in the European honeybee population. The pileup files were interpreted by the PoPoolation2 utility mpileup2sync (Kofler et al., 2011) for the Sanger Fastq format, with a minimum quality of 20 and were finally converted to allele counts and sequencing depth files using a custom-made script. In addition, for each of these 34 colonies 4 male offspring of the queen, genetically equivalent to queen gametes, were individually sequenced as in Wragg et al. (2021) (Supplementary Table ST4). In order to reduce computation time this analysis was performed on a subset of about 50000 markers. These markers were selected following the criteria: 1) maximum of two polymorphic sites within a 100 base pair window, 2) only one representative marker per linkage disequilibrium block with *r*^2^ higher than 0.8, 3) variance between allele frequencies in the different genetic backgrounds higher than zero, to allow for population identification and 4) sampled so that the minor allele frequency follows a uniform distribution. This selection led to exactly 48 334 markers in the experimental dataset.

#### Genetic ancestry

For each colony, using pooled sequencing data, the queen genetic ancestry *q^c^* was estimated using AM as described above. For the male offspring data, for each colony two ways to estimate the genetic ancestry were considered:

1. By averaging the genetic ancestry vectors of the four males as estimated by ADMIXTURE.
2. By first reconstructing the queen genotype from the male offspring data (see below) and then analysing the resulting genotype with ADMIXTURE.

#### Genotype reconstruction

For pool sequencing data, queen genotypes were reconstructed using HP, considering the 34 colonies jointly. For the male offspring data, queen genotypes were reconstructed by first estimating the genotype probabilities at each locus from individual data at the four individually sequenced male offspring. Our goal is to reconstruct the genotype of a parent at a locus (*g_l_*) (here the queen) from the haploid genotypes of a set of *n_g_* gametes (here the male offspring). Let *R* be the random variable of the number of reference alleles observed in the offspring and assume that there is a per allele sequencing error equal to *ϵ*, then the genotype likelihoods can be computed from the sampling distributions:

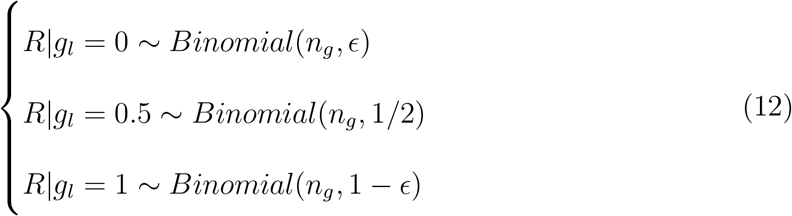

To compute the genotype posterior probability when *r_l_* reference alleles are observed at a locus, we specify a uniform prior on the three possible genotypes, so that 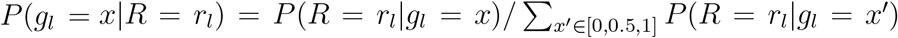. For our application, we fixed *ϵ* = 10^−3^ and *n_g_* is four as described above. Because we have only four drones per colony in this dataset, there is still some uncertainty in the genotype of the queen. For example the highest posterior probability achievable for a genotype with *n_g_* = 4 is ≈ 0.94. This has to be taken into account when comparing the genotypes reconstructed from the offspring data and from the pool sequencing data: the concordance between the two approaches has to be measured with respect to what is expected between the true genotype of the queen and the one reconstructed from noisy data (either offspring or pool sequencing). Unfortunately we do not know the true genotype of the queen in our dataset but we can measure the concordance between the genotype reconstructed with four male offspring to the true genotype of the queen using data from Liu et al. (2015). In this dataset, genotypes of 13 to 15 offspring are available for three colonies. With that many offspring the genotype of the queen can be reconstructed with certainty and be compared to the one obtained by downsampling the data to four offspring per colony. Therefore, for each of the three colonies in Liu et al. (2015), we called the offspring genotypes at the set of markers present in the diversity panel, reconstructed the queen genotype using (i) all offspring (*n_g_* = 15 or 13) and (ii) a 100 randomly downsampled datasets consisting of four offspring only.

## Results

In this study we developed statistical models to estimate genetic ancestry and queen genotypes from pool sequencing data from workers of the colony. Simulations, from independent and linked markers, were performed to evaluate the performance of our models in terms of queen genetic ancestry inference and genotype reconstruction. The scenarios are described in Figure S1. Moreover, these models were applied to an experimental dataset composed of both pool sequenced data and individual male offspring of the queen. In fact male offspring of the queen, haploid individuals coming from unfertilised queen gametes, are direct sampling of the queen genetics and their use is often suggested in literature as a proxy for queen information.

### Validation on simulations

#### Genetic ancestry

For independent markers, correlations between simulated genetic ancestries and estimated genetic ancestries using the Admixture Model (AM) ranged between 0.88 and 0.9 depending on the genetic background and for linked markers correlations between genetic ancestries estimated using ADMIXTURE (Alexander et al., 2009) on the queen genotypes simulated from real data and estimated by AM ranged between 0.93 and 0.95 depending on the genetic background (Figure 1). In addition to the 15 scenarios listed we also estimated genetic ancestries by AM on scenarios in which queen and drones had divergent ancestries (Supplementary table ST1). We observed that shifting from the initial hypothesis that queen and drones come from the same origin led to highly biased genetic ancestry estimations with AM (Figure S3). It should be noted that the statistical model under AM is based on the assumption that markers are independent. To match this assumption a subset of 1000 markers, rather than the whole genome, was used to estimate genetic ancestry for simulations with linked markers. These results show that AM outputs accurate genetic ancestry estimates and show inference with high agreement to standard population genetics models such as ADMIXTURE, under the assumption that queen and drones are of the same origin. Moreover, the observed results confirm that using only a subset of ancestry informative markers, here 1000 from the whole genome, is sufficient to accurately estimate genetic ancestries using AM.

**Figure 1:**
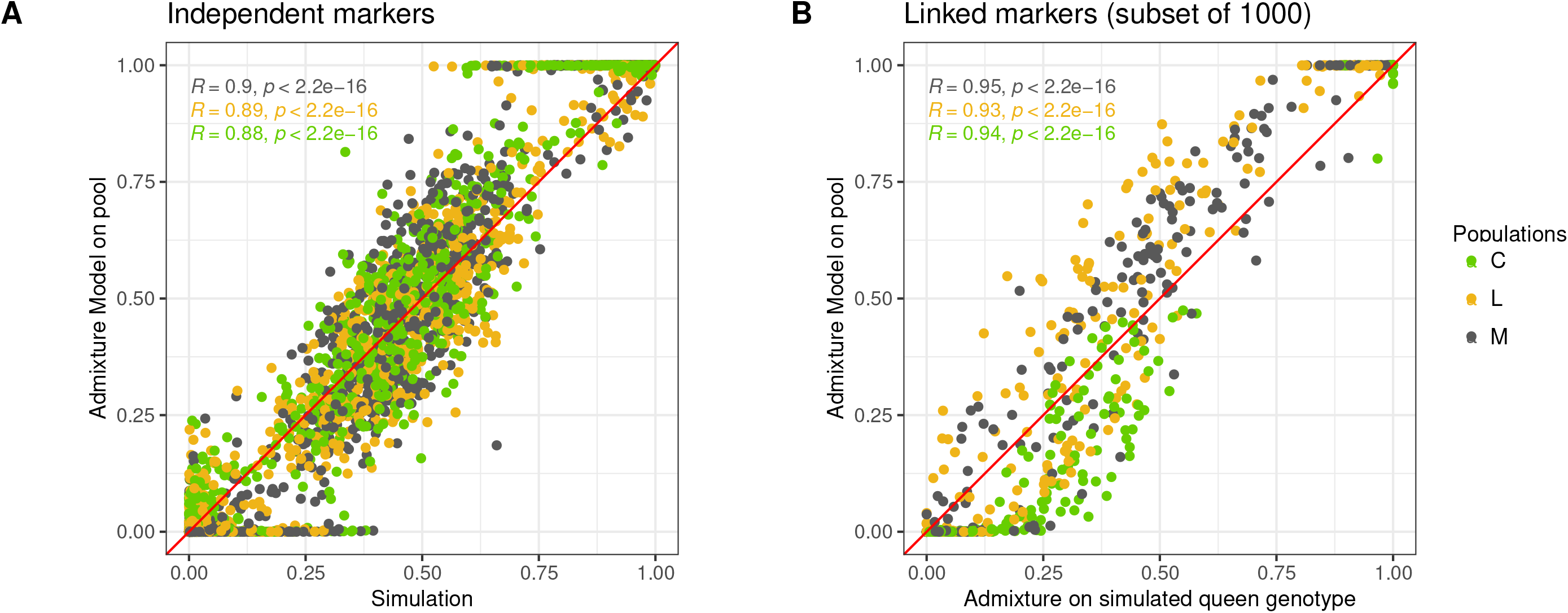
Genetic ancestry comparison. Regression of the genetic ancestry vectors estimated by the Admixture Model against simulated. Genetic ancestries estimated with AM against simulated for independent markers, for each scenario, each colony, for each genetic background (15 * 100 * 3) (A) or estimated with AM against by ADMIXTURE for simulations for linked markers (subset of 1000), for each scenario, each colony, for each genetic background (15 * 20 * 3, the number of simulated colonies is lower due to limitation in the number of individuals to sample from in the real dataset) (B). The red line represents the regression with intercept 0 and slope 1, meaning perfect agreement between the two estimates. Values for spearman rank correlations between ancestry vectors are shown in the top left corner for each of the three genetic backgrounds in green *Apis m. caucasia* C, in yellow *Apis m. ligustica & carnica* L and in grey *Apis m. mellifera* M.

#### Genotype reconstruction

One major assumption of the Homogeneous Population Model (HP) is that colonies within the population are of homogeneous genetic ancestries. Therefore, using simulations for linked markers across the whole genome, we compared and clustered all the simulated colonies based on their genetic ancestries estimated by AM. In our study we assume that colonies come from a mixture of three main genetic backgrounds (as described in Wragg et al. (2021)), we thus clustered our simulated colonies in seven groups from pure to hybrid genetic types (Figure 2).

**Figure 2:**
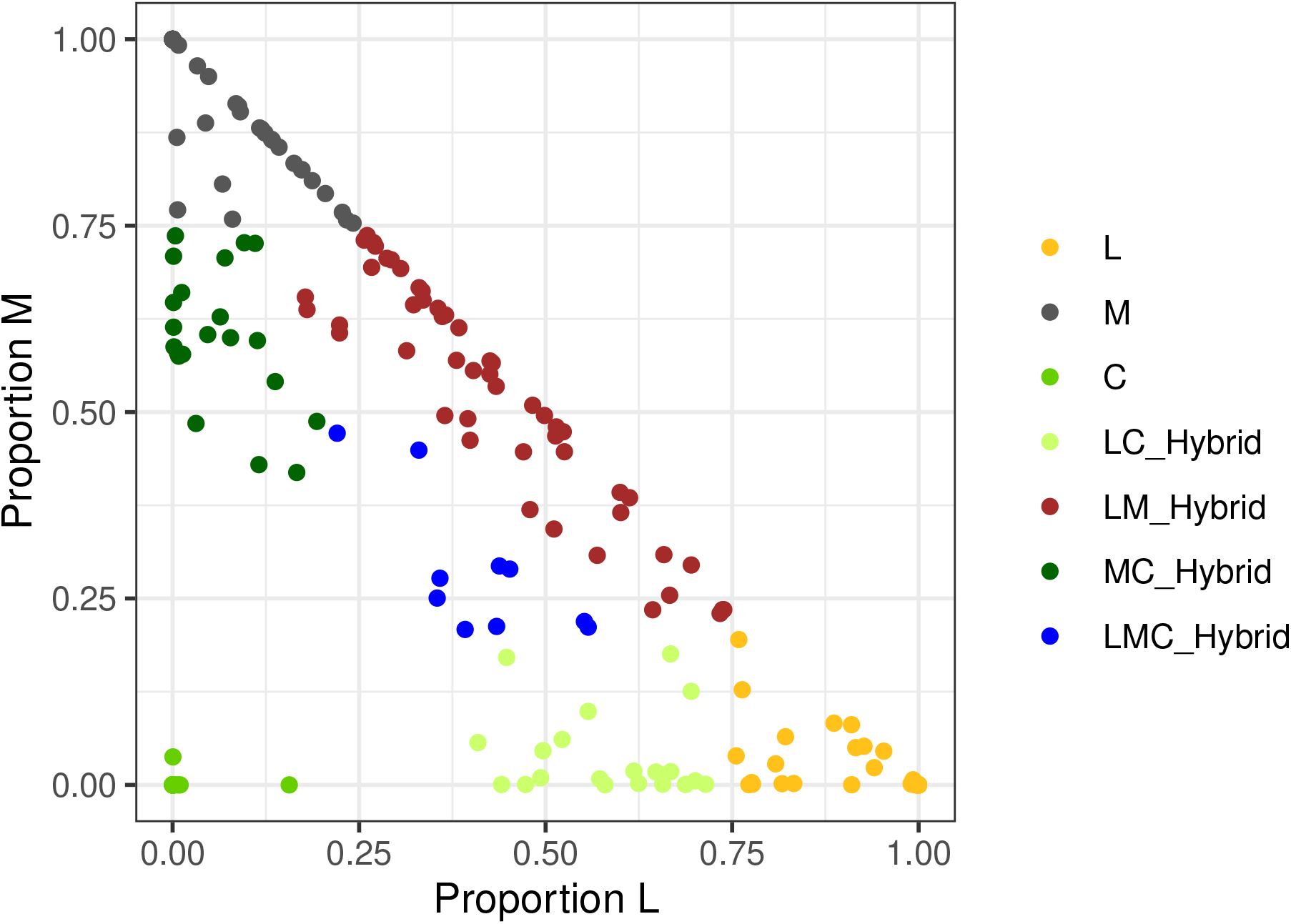
Genetic ancestries for the simulated colonies as estimated by the Admixture Model. Two dimensions plot of genetic ancestries estimated by AM for colonies simulated for linked markers. X and y axis give the genetic ancestry values in two of the three populations of honeybee in our dataset, for all the colonies in all scenarios (20 * 15) simulated for linked markers after estimation of their genetic ancestry vectors by the AM model. Individuals can be grouped by genetic ancestry. Here we decided on seven groups, each in a different colour, in yellow *Apis m. ligustica* + *Apis m. carnica* L, in grey *Apis m. mellifera* M, in green *Apis m. caucasia* C, in light green hybrids *Apis m. ligustica* and *Apis m. caucasia*, in brown hybrids *Apis m. ligustica* + *Apis m. carnica* and *Apis m. mellifera*, in dark green hybrids *Apis m. mellifera* and *Apis m. caucasia* and in blue the three ways hybrids.

Thereafter, to evaluate queen genotype reconstruction performance we implemented the Homogeneous Population Model (HP) on our seven groups of homogeneous colonies for linked markers. As the HP model does not make the assumption of independence of markers the inference could be performed on the whole genome, approximately 7 million markers. Across all simulations and all scenarios, we observed a good correlation between the rate of genotype agreement between simulated and estimated genotypes and the associated estimated genotype probability. In other terms genotypes inferred with a high probability are often correctly predicted by HP whereas genotypes inferred with a low probability are often wrongly predicted by HP, making genotypes with a probability close to 0.5 the hardest to infer precisely. The calibration of the HP model for genotype reconstruction, measured as the Area under the Curve between agreement rates and probabilities was 0.055 (Figure 3A), when AUC ranged between 0, for perfect correlation and 0.5 for completely imperfect correlation. A large proportion of the markers have probabilities close to zero or to one, making the genotypes drawn for these markers close to certain (Figure 3A). As expected we observed that the genotyping error rate decreases slightly when the best genotype probability threshold increases meaning that filtering for markers with higher best genotype probability leads to more accurate genotype reconstruction. However such filtering is accompanied with a small reduction in genotype call rate. For example if no filtering on best genotype probability is applied, 100% of the genome will be reconstructed with an average genotyping error rate of 4%, if filtering for markers with best genotype probabilities above 0.9 is applied about 95% of the whole genome will be reconstructed with an average genotyping error rate as little as 2% (Figure 3B). Additionally we observed that the genotyping error rate increased when the MAF threshold increased meaning that filtering on MAF might cause an increase in genotyping error, accompanied by a drastic reduction in genotype call rate (Figure 3C). Minor Allele Frequency and best genotype probability are highly linked as markers with low MAF tend to be easier to infer with high probability. In our simulation a large proportion, more than 50%, of the whole genome is composed by markers with MAF below 0.05. Yet applying a filter on best genotype probability does not seem to highly impact the distribution of MAF on the whole genome (Figure S4). Rather than filtering on MAF we suggest to filter on best genotype probability, for example equal to or greater than 0.95. Indeed, such filtering will improve the queen genotype reconstruction accuracy without heavily impacting the allele frequency distribution of the markers genotyped on the whole genome. In fact, we observed that genotyping error, on the whole genome and without filtering, is on average about 3% (Figure 3D). After applying a filter on best genotype probability equal to or greater than 0.95 genotyping error becomes on average as low as about 2%.

**Figure 3:**
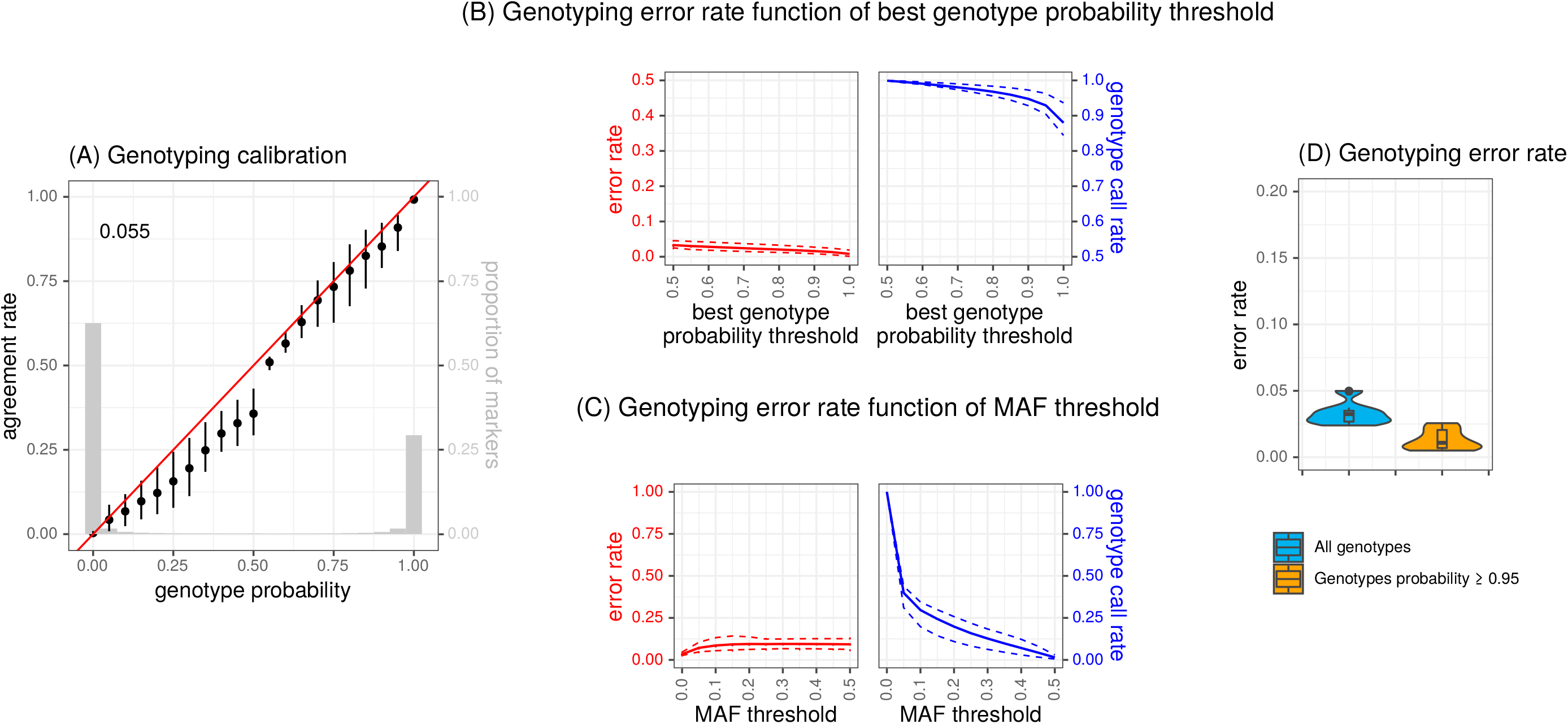
Queen genotype reconstruction. For linked marker simulations on the whole genome, values averaged across all colonies and scenarios. A) Genotyping calibration, each point represents the genotype agreement rate per genotype probability value with bars representing the quantiles 95% to 5%. The red line is the regression with intercept 0 and slope 1. Value for Area Under the Curve between perfect and observed calibration is shown in the top left corner. The grey histogram represents the proportions of markers in each bin of genotype probability. B) Genotyping error rate as function of best genotype probability threshold. In red, the solid line represents the average genotyping error rate across all scenarios as a function of the best genotype probability, the dotted lines are the quantiles 95% and 5%. In blue, the solid line represents the average genotype call rate, across all scenarios, if thresholds were applied on the best genotype probability, the dotted lines are the quantiles 95% and 5% for genotype call rate. C) Genotyping error rate as function of Minor Allele Frequency threshold. As for B), in red, the solid line represents the average genotyping error rate across all scenarios as a function of the MAF threshold, the dotted lines are the quantiles 95% and 5% for genotyping error rate. In blue, the solid line represents the average genotype call rate, across all scenarios, as a function of the MAF threshold, the dotted lines are the quantiles 95% and 5% for genotype call rate. D) Violin plot of the genotyping error, for all markers or filtering on best genotypes probability equal to or greater than 0.95.

These results show average estimates across all simulation scenarios and colonies after grouping based on genetic ancestry. Detailed results for calibration and genotyping error are presented Figure S5.

To conclude, using simulations we confirm that the statistical model AM performs similarly to ADMIXTURE leading to highly accurate genetic ancestry inference. A small set of markers, as low as 1000 in our example where genetic background differentiation is strong, seems sufficient to accurately estimate genetic ancestry with AM. Using simulation of linked markers across the whole genome we confirmed that HP reconstructed queen genotypes with high accuracy. Furthermore, we inferred the impact of MAF and best genotype probability thresholds on the genotype call rate and the associated genotyping error rates, giving the advice to filter on best genotype probability equal to or greater than 0.95 to reduce genotyping error, without drastic loss of predicted markers and while preserving allele frequency distribution across the genome.

### Application on experimental data

To further evaluate the performance of the AM and HP models, we analyzed real data on honeybee colonies for which 4 drones were individually sequenced (see Materials and methods).

#### Genetic ancestry

For each colony, the genetic ancestry of the queen was estimated either from the group of male offspring or from the pool sequences of workers. Genetic ancestry from worker pool sequence were estimated using the Admixture Model (AM). For male offspring, it was estimated with ADMIXTURE (Alexander et al., 2009) either using the male offspring directly (admix_males) or from the genotype of the queen reconstructed using male offspring (admix_proba), as described in the Material and Methods section. Using male offspring data directly (admix_males) or through queen genotype reconstruction (admix_proba) genetic ancestry from ADMIXTURE were virtually equal with a Mean Squared Difference (MSD) of 1.4 × 10^−3^ (standard deviation 1.1 × 10^−3^). Comparing estimates based on male offspring versus worker pool sequence (AM) MSD were slightly higher with 0.024 and 0.026 with standard errors of 0.025 and 0.021 for admix_males and admix_proba respectively (Table 2). Out of the 34 experimental colonies most of the genetic ancestry estimated using queen reconstructed genotypes from worker pool sequencing data, male offspring or using individual sequencing of male offspring gave nearly identical q vectors (Figure S6).

**Table 2:** Genetic ancestry Mean Squared Difference between data and models.

|  | queen from males<br>vs males | queen from pool<br>vs males | queen from pool<br>vs queen from males |
| --- | --- | --- | --- |
| **model_i** | admix_proba | AM | AM |
| **model_j** | admix_males | admix_males | admix_proba |
| **min** | 1.36E-05 | 2.94E-04 | 1.35E-03 |
| **mean** | 1.43E-03 | 0.024 | 0.026 |
| **median** | 1.15E-03 | 0.014 | 0.020 |
| **max** | 4.19E-03 | 0.085 | 0.082 |
| **sd** | 1.16E-03 | 0.025 | 0.021 |

Genetic ancestry Mean Squared Differences for different data and models on experimental colonies. Minimum, average, median, maximum and standard deviations are calculated for each combination.

#### Genotype reconstruction

To validate queen genotype reconstruction from worker pool sequence on our experimental dataset we used publicly available data from Liu et al. (2015) on three colonies for which both queen and 13 to 15 male offspring were individually sequenced were used. Of the 50000 selected markers only 14988 were available, as polymorphic SNPs, on the dataset from Liu et al. (2015). This reduction in the number of markers available for the analysis can be explained as the population used for SNP calling was composed of fewer individuals from a unique and uniform origin in the dataset from Liu et al. (2015). We compared queen genotypes reconstructed from worker pool sequence and queen genotype reconstructed on probabilities from four male offspring (pool/offspring) on the experimental dataset, genotypes from individually sequenced queens and queen genotype reconstructed on probabilities from four male offspring (queen/offspring) and pairs of queen genotype reconstructed on probabilities from four independent male offspring (offspring/offspring) on the dataset from Liu et al. (2015). Genotype concordance was on average 0.94 (standard deviation 0.03), 0.96 (standard deviation 0.01) and 0.92 (standard deviation 0.01) for pool/offspring, queen/offspring and offspring/offspring respectively (Figure 4). The highest concordance is observed between the actual queen genotypes and its reconstruction from four male offspring; however queen genotype reconstruction from pool and from male offspring seem to present similar concordance than when pairs of independent male offspring are compared. The few colonies showing more discrepancy between genetic ancestry estimates always showed a genetic ancestry from worker pool sequence mostly divergent from the estimates based on males, despite having high concordance between genotype reconstruction. This can be either due to limitations in AM when it comes to disentangling queen genotype from cohort of inseminating drones in the worker pool sequencing data, to the fact that sampling only four male offspring is not sufficient to accurately represent the queen genetic ancestry, because of genetic con-tradiction between the queen that produced the male offsprings and the one that produced the workers or to a biais in the markers used for AM. However, this validation confirms that queen genotype reconstructed using worker pool sequencing data performs as well as individually sequencing multiple male offspring. Additionally we showed, on the data from Liu et al. (2015), that increasing the number of male offspring individually sequenced to six, eight or even ten improved the genotype concordance quite substantially (Figure S7) with eight and ten male offspring showing a concordance between reconstructed and real genotype close to one.

**Figure 4:**
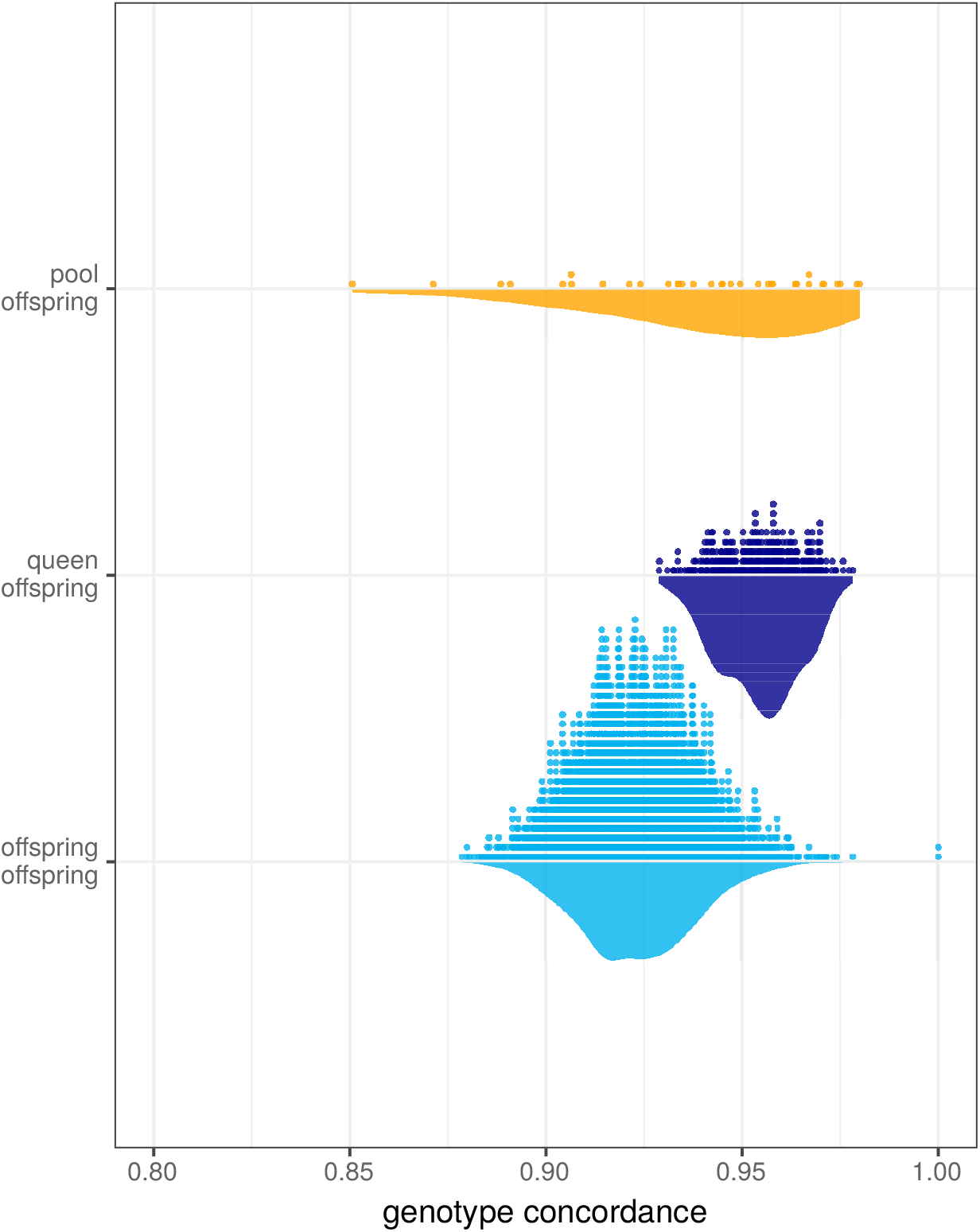
Concordance between queen genotype reconstruction based on different data. Concordance between reconstructed genotypes from different data types. The densities, bottom, represent the concordance, only for markers after filtering for best genotype probability equal to or greater than 0.94, between i) queen genotype reconstructed from pool sequencing data using HP and queen genotype reconstructed from genotype probabilities (pool/offspring), based on four male offspring for experimental colonies, in orange ii) queen genotype reconstructed from genotype probabilities based on four male offspring for a 100 sampling events and actual queen genotypes from the Liu et al. (2015) (queen/offspring), in dark blue and iii) pairs of queen genotype reconstructed from genotype probabilities based on four male offspring for independent sets of individuals with the data from Liu et al. (2015) (offspring/offspring), in light blue. Concordance values for each test are represented as dots, top, and as density distribution, bottom.

To summarise, the difference between genetic ancestry estimated from male offspring or worker pool sequencing data, using AM, were small. Queen genotype reconstruction from worker pool sequencing data was in agreement with queen genotype reconstructed from male offspring. This value was slightly lower than when comparing queen reconstructed genotypes from male offspring with the real queen genotype and slightly higher than when comparing queen reconstructed genotypes from different sets of male offspring of the same queen. HP on worker pool sequencing data is an accurate alternative to individually sequence a limited number of male offspring of the queen when one wants to access the queen genotype.

## Discussion

The past decade has seen the growth of the molecular genomics era with the development of new sequencing platforms and technologies, one of them being pool sequencing. This technology allows for the combination of multiple individuals in one sequencing experiment, reducing drastically preparation and sequencing costs and therefore making high depth sequencing available for a wide variety of samples. Traditionally pool sequencing is used to perform analysis on multiple individuals from a population. Additionally pool sequencing might be of interest when group level information is desired as for example in the context of eusocial organisms. In such cases the pool will represent a meta-individual of the colony rather than a population. One pitfall of using such sequencing method it that the outcome of pool sequencing comes in the form of allele read counts and sequencing depths rather than diploid genotype observations making it a non standard format for downstream analysis.

So far only a few programs, for example Popoolation (Kofler et al., 2011) and CRISP (Bansal, 2010) for SNP calling, Plink (Purcell et al., 2007; Chang et al., 2015) and the R package poolfstat (Hivert, Leblois, Petit, Gautier, and Vitalis, 2018; Gautier, Vitalis, Flori, and Estoup, 2021) for population genetics or GEMMA (Zhou and Stephens, 2012) and LDAK (Speed et al., 2020) for association study handle non genotype data. However, when considering eusocial insects from the same colony as a pool we might break underlying assumptions made by these models. In fact, eusocial insects present characteristics deviating from what could be expected in a standard population used for pool sequencing experiments. First, in hymenopterans, reproductive systems are often polyandric, leading to non standard genetic relationships across individuals in the colony. Second, traits of interest are likely to be measured at the colony level. Therefore, in order to avoid computational limitations and biases that could be brought by the use of pool sequencing with unadapted models one may want to infer individual genetic information (e.g. ancestry and genotypes) from a pool from the group. In honeybee, for instance, a colony can be considered as a polyploid organism (with two major chromosomes, coming from the queen and being present in the whole population, and about 15, the number of inseminating drones, minor chromosomes) constituted of haploid male offspring of the queen that can be described as ‘’flying gametes’‘ as they come from queen unfertilised eggs and diploid female offspring of the queen, worker bees, descendant from the mating of a queen with a cohort of about 15 inseminating drones. Genetic relationships between colony inmates is more complex than in other animal species as they range between 1 to 0.25 depending on the patriline from which the individual belongs (Oxley and Oldroyd, 2010). The honeybee queen carries the largest part of the genetic information of the colony and is the producing organ of the next generation making it a favored pathway for breeding selection. In addition, the honeybee populations used by breeders and beekeepers are often highly structured with vast differences between genetically pure and highly admixed colonies. The honeybee population has been influenced by domestication and selection performed by beekeepers often on traits measured at the colony level making the use of pool highly relevant. These features make the use of *Apis mellifera* as a model organism, to develop statistical models to use pool sequencing data, greatly relevant. Moreover we also benefit from the available knowledge on the organism compared to other eusocial insects. For example we can exploit the diversity panels, such as built in Wragg et al. (2021), as priors in our models to facilitate inference. In this context the developed methods are expected to be easily applicable to organisms with lower level of population stratification, as can be for some other eusocial insects.

Here we present two statistical models to infer queen information from pool experiment data. First, the Admixture Model (AM) allows to infer queen genetic ancestry from worker pool sequencing data knowing expected allele frequencies in a reference populations with high correlation between predicted and expected ancestry (about 0.9) and computational efficiency as it can be run rapidly for each colony independently, thus parallelisable, on a small subset of markers. Second, the Homogeneous Population Model (HP) allows for an accurate queen genotype reconstruction with as little as 2% genotyping error. This model takes advantage of the information from other colonies of the group to complete genotype reconstruction, making the assumption that colonies within a group are of homogeneous genetic ancestry. Within the context of population genetics study, when genetic ancestry is unknown prior to the analysis and knowing the results of this study we suggest to first infer genetic ancestry using AM for all the colony DNA pools of interest, then group them based on similarities in their ancestries and perform genotype reconstruction on these groups separately with HP. Therefore, we propose to use our statistical models sequentially to reach highly accurate genotype reconstruction. To date a common way to infer honeybee queen genotype without manipulating and sacrificing this queen is to perform pool sequencing on multiple honeybee queen male offspring (Petersen et al., 2020). For this purpose Jones et al. (2020) suggests, using theoretical estimations, to sequence at least 30 individuals. This procedure requires to be able to identify and sample enough male offspring from the colony, which is not always easy depending on the season, the colony and the time available for sampling. An alternative is to individually sequence multiple honeybee queen male offspring, in such case, the number of individual sequences is the limiting factor to an accurate queen genotype reconstruction with at least eight to ten individuals needed to accurately deduce queen genome phase, that we cannot obtain from a pool experiment, and to lower the risk of incorrect genotype reconstruction (Figure S7). Using real data we saw that our statistical models, based on pool sequence experiments, reconstructed queen genotypes at least as well as using four individual male offspring sequences. Queen genotype reconstruction from pool sequencing data from workers of the colony appears to be a relevant alternative, cheaper as only one sequencing procedure needs to be performed. Simulations, of independent and linked markers, and the experimental field dataset concluded that we could estimate honeybee queen genetic ancestry and genotype accurately and efficiently using our methods.

Despite the efficiency of the statistical models described in this study some limitations have been identified and further improvements can be conducted. One crucial assumption of our model is that honeybee queens and inseminating drones have similar genetic ancestry, which is often true when natural breeding is conducted. However this assumption might be broken when conducting queen artificial insemination for breeding purposes, in extremely controlled breeding environments or even when the breeding environment is ’polluted’ by unexpected genetics. In fact, when queen and inseminating drones have highly divergent ancestries our models will estimate biased genetic ancestry and queen genotypes (Figure S3). Additional external information is necessary to account for heterogeneity in the origin of breeding parents of the pool. One way to do so would be by implementing a two step reconstruction algorithm focusing first on the inseminating drones allele frequencies, for example using information on the breeding practices or sampling drones from the environment as a representation of the mating cohort. Once information on the mating cohort is available it can be easily implemented in our model by adapting the prior in the equation (6). In this study we performed simulations of pool experiments with a sequencing depth of 30x. In practice, and especially in the context of non-model organisms, such sequencing depth might be difficult to reach either due to sequencing cost or to genetic material availability. Therefore, we also tested the simulations with a depth of 10 or 100. We compared our results in terms of genotyping error rate and genotype call rate on the genome after filtering for best genotype probability. In Figure S8 we can see that increasing sequencing depth from 10 to 30 improved the accuracy of genotype inference and the genotype call rate. At high sequencing depth, 100, we observed higher genotyping error rate overall and limited improvement in the fraction of markers inferred with certainty. It is likely that some level of heterogeneity within the groups used to reconstruct queen genotype led to wrong decisions at higher sequencing depth. Increasing sequencing depth seems to cause higher sensitivity to the hypothesis of homogeneous population by the statistical HP model. One option to reduce this impact would be by grouping colonies based on their genetic ancestries to a more refined scale. Indeed, further developments in the HP model could allow one to take into account a level of heterogeneity in the population to reduce the sensitivity of the model to the homogeneity assumption.

We observed that HP performed better, had a lower genotyping error rate, if inferred genotypes along the genome were filtered based on their certainty, measured as a probability. In our simulations such filtering did not affect the allele frequency distribution and reduced only slightly the number of inferred markers along the genome while reducing genotyping error rate (Figure S4). An imputation step would contribute to the improvement of genome reconstruction completeness. Also taking into consideration Linkage Disequilibrium (LD) along the genome to refine the genotypes inferred by HP could be adapted in our statistical model. Such development would benefit from identification of haplotype blocks in the honeybee genome (Saelao et al., 2020; Wallberg, Schöning, Webster, and Hasselmann, 2017; Wragg et al., 2016; Wragg et al., 2021) tagging the different *Apis mellifera* populations. An efficient strategy would be to reconstruct queen genotypes with HP, filter on genotype probability to retain only markers from which reconstruction is satisfying and then apply an imputation step taking into account known haplotype blocks and LD between markers.

To conclude, colony pool sequencing data can be used to infer queen genetic ancestry when knowing allele frequencies in reference populations present in the environment. Moreover, using pool sequencing data across multiple colonies of homogeneous genetic ancestry in which queen and inseminating drones come from a similar origin, it is possible to reconstruct honeybee queen genotypes accurately. Such genotypes are valuable for example to run population genetics analysis and association studies with mainstream models currently available and genetic ancestry estimates can be useful for selective breeding purposes. Additional developments to take into consideration some level of heterogeneity, discrepancy of origins between queen and inseminating drone cohort and linkage disequilibrium along the genome will help further increase genotype reconstruction accuracy. The statistical models described in the study have been designed within the context of eusocial hymenoptera but tested solely on *Apis mellifera*. Such models could be tested within the framework of studies on other eusocial species with multiple mating of a single queen (Micheletti and Narum, 2018) and with known genetic diversity panels to estimate priors for allelic frequencies.

## Data accessibility statement

Scripts developped to perform the simulation are available at xxxxx for download. The vcf file containing the filtered SNPs and the complete diversity panel can be found in Wragg et al. (2021). The list of 628 individuals used in this study as well as the list of reference individuals and individuals (male offsprings) used for validation can be found in the Supplementary Table S1, together with their accession names. The pool sequencing experiment data for the 34 colonies used for validation can be found at xxx. The external data set used for validation can be found in Liu et al. (2015).

## Author’s contributions

AV, BS, FM, BB, YLC and AD designed the data collection. FM, BB and YLC performed the data collection. KT, and EL performed the laboratory preparation of the samples, DNA extraction, library preparation and sequencing. SEE, BS and AV designed the study. BS developed the methods and wrote the models. SEE designed and performed the simulations and model comparisons. SEE, BS and AV interpreted the results. FM, YLC, LG, and AD contributed to the discussion. SEE, BS and AV drafted and reviewed the manuscript. All authors have read and approved the manuscript.

## Acknowledgements

This study was performed with the support of the ITSAP team for the maintenance of the honeybee colonies and the data collection, the sequencing platform GeT-PlaGe, Toulouse (France), a partner of the National Infrastructure France Génomique, thanks to support by the Commissariat aux Grands Invetissements (ANR-10-INBS-0009), for the sequencing and especially Olivier Bouchez. Bioinformatics analyses were performed on the computing facility Genotoul. This research was funded by the Ministère de l’Agriculture de l’Agroalimentaire et de la Forêt within the framework of MOSAR RT 2015-776 project and the Ministère de l’Agriculture de l’Agroalimentaire et de la Forêt and Investissement d’avenir for BeeStrong PIA P3A project. Thanks to Claude Chevalet for the initial discussions on the idea, the members of the BeeStrong project, Florence Phocas and François Guillaume, for their contributions to the discussion during the development of this study.

## Supplementary Figures

**Figure S1:**
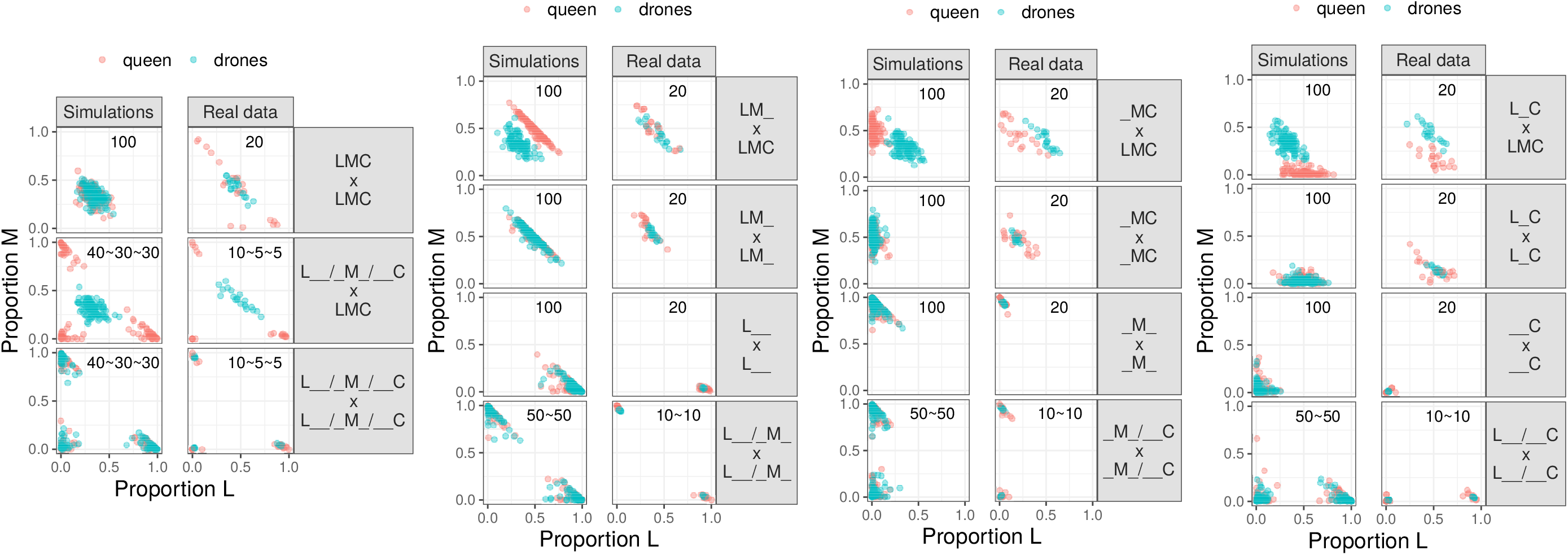
Simulated genetic ancestries for queen and drones. Two dimensions plot of genetic ancestries simulated for queens (pink) and drones (blue) for all scenarios.

**Figure S2:**
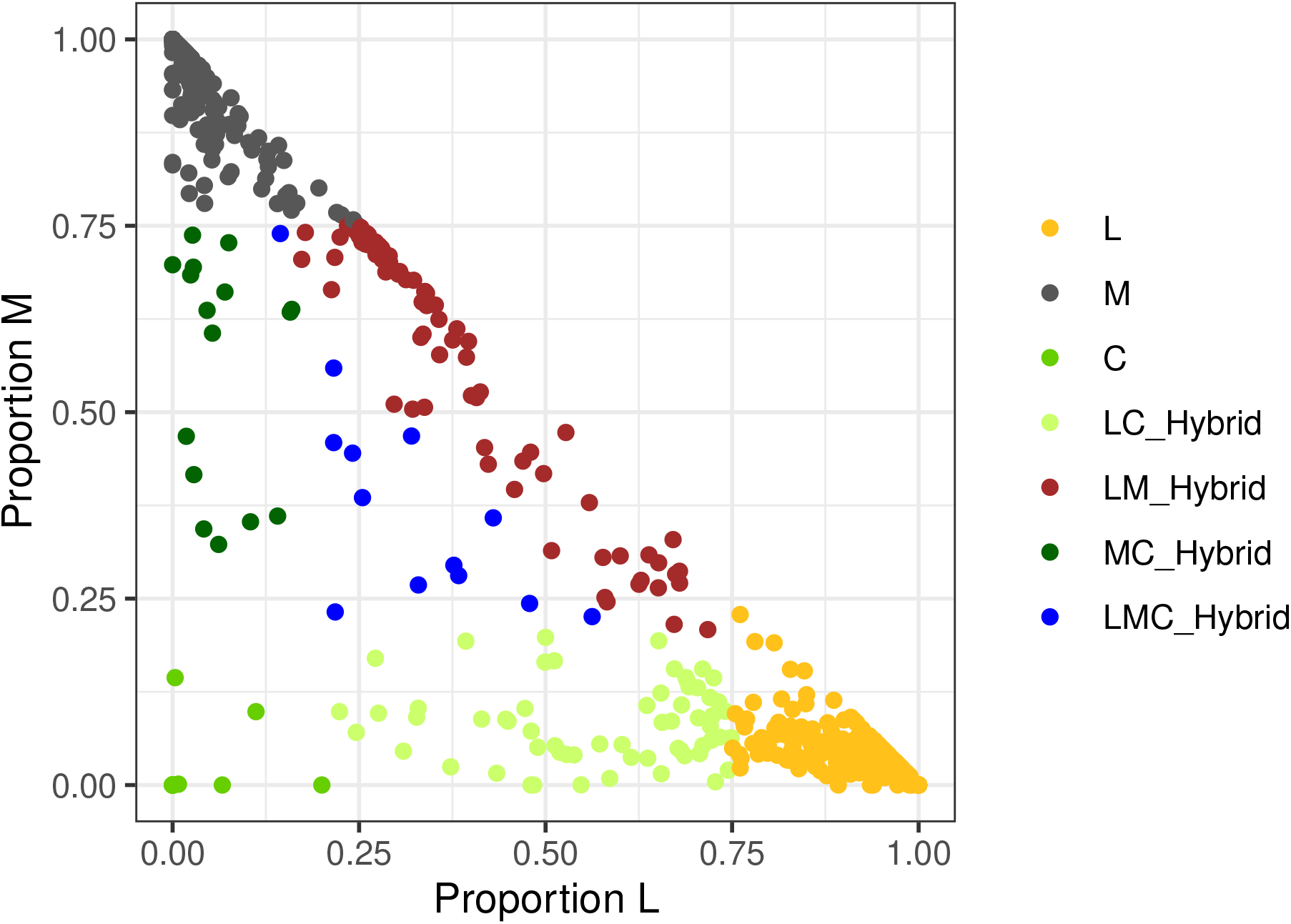
Genetic ancestries of 628 male individuals from the diversity panel of Wragg et al. (2021) Two dimensions plot of genetic ancestries for the individuals from the diversity panel. Individuals can be grouped by genetic ancestry. Here we decided on seven groups, each in a different colour, in yellow *Apis m. ligustica* + *Apis m. carnica* L, in grey *Apis m. mellifera* M, in green *Apis m. caucasia* C, in light green hybrids *Apis m. ligustica* and *Apis m. caucasia*, in brown hybrids *Apis m. ligustica* + *Apis m. carnica* and *Apis m. mellifera*, in dark green hybrids *Apis m. mellifera* and *Apis m. caucasia* and in blue the three ways hybrids.

**Figure S3:**
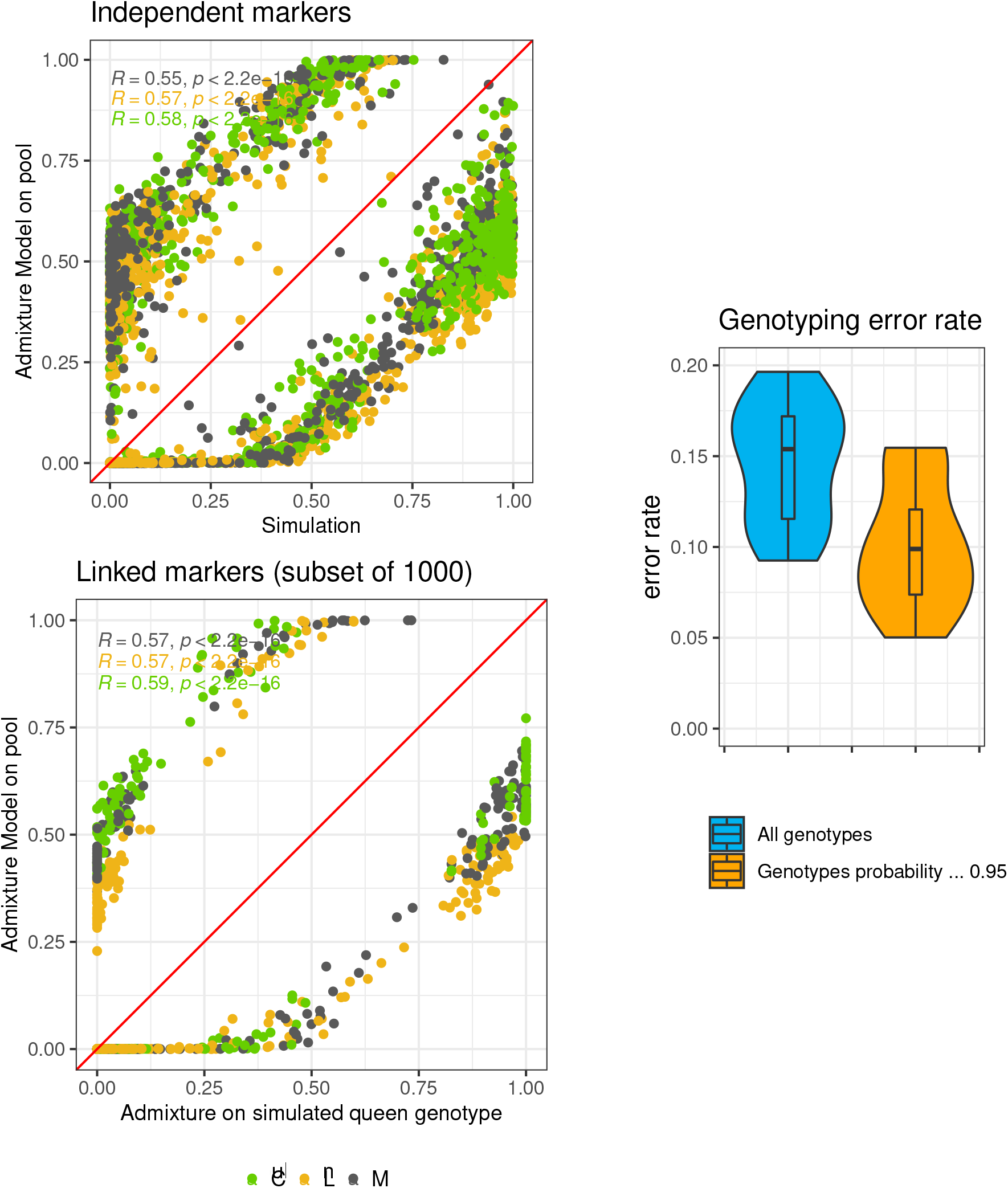
Genetic composition and genotyping error when queen and drones come from different ancestries. Detailed information are available in Supplementary Table ST1

**Figure S4:**
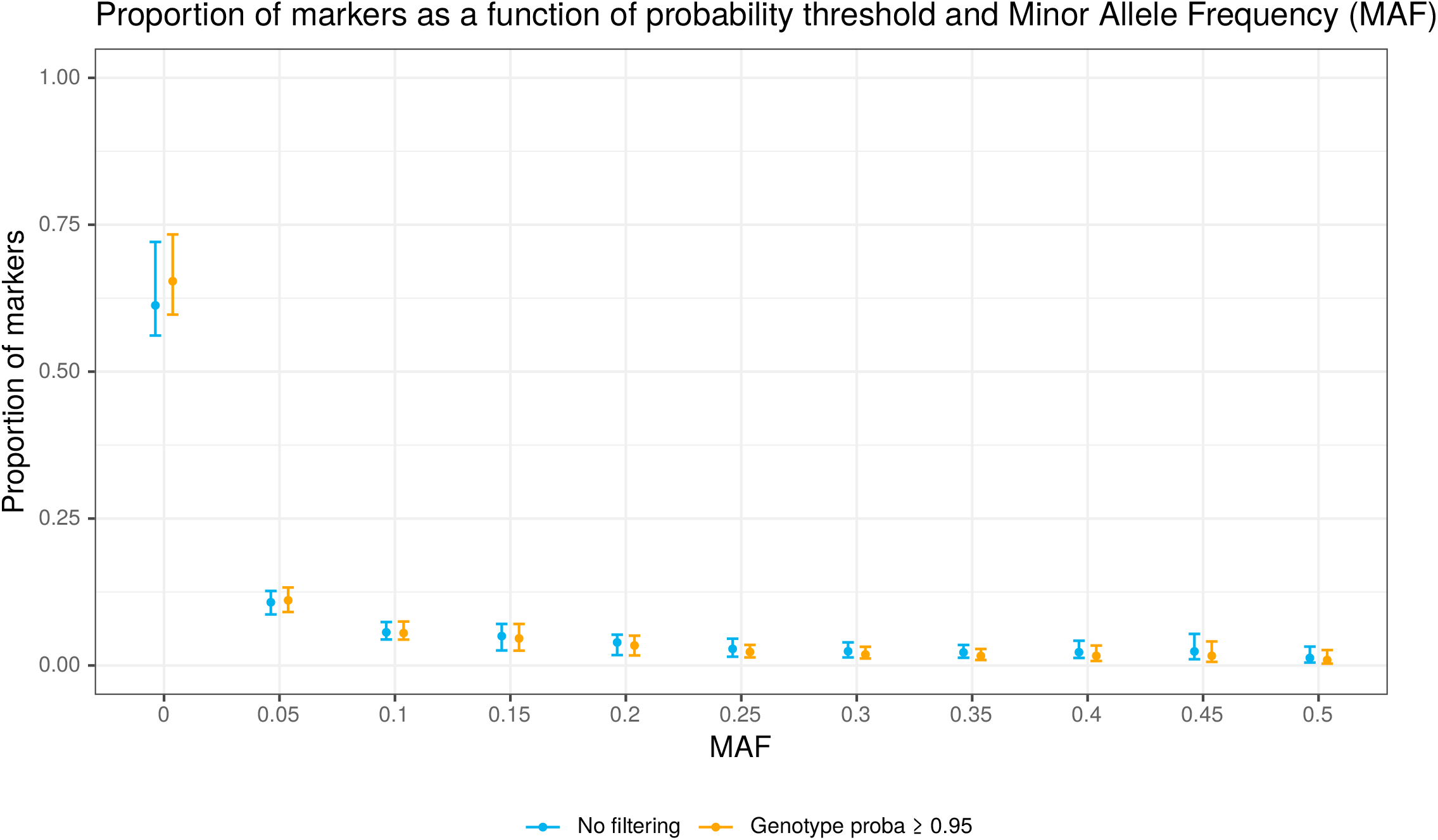
Proportion of markers in each MAF categories under best genotype probability thresholds. Representation for each MAF category of the mean, with interval, proportion of markers. For MAF 0 to 0.5 we represented the mean and the quantiles 95% and 5% proportion of markers across all simulations. In blue without filtering on best genotype probability, in orange after filtering for markers with best genotype probability equal to or greater than 0.95 on the whole genome or only on these filtered markers.

**Figure S5:**
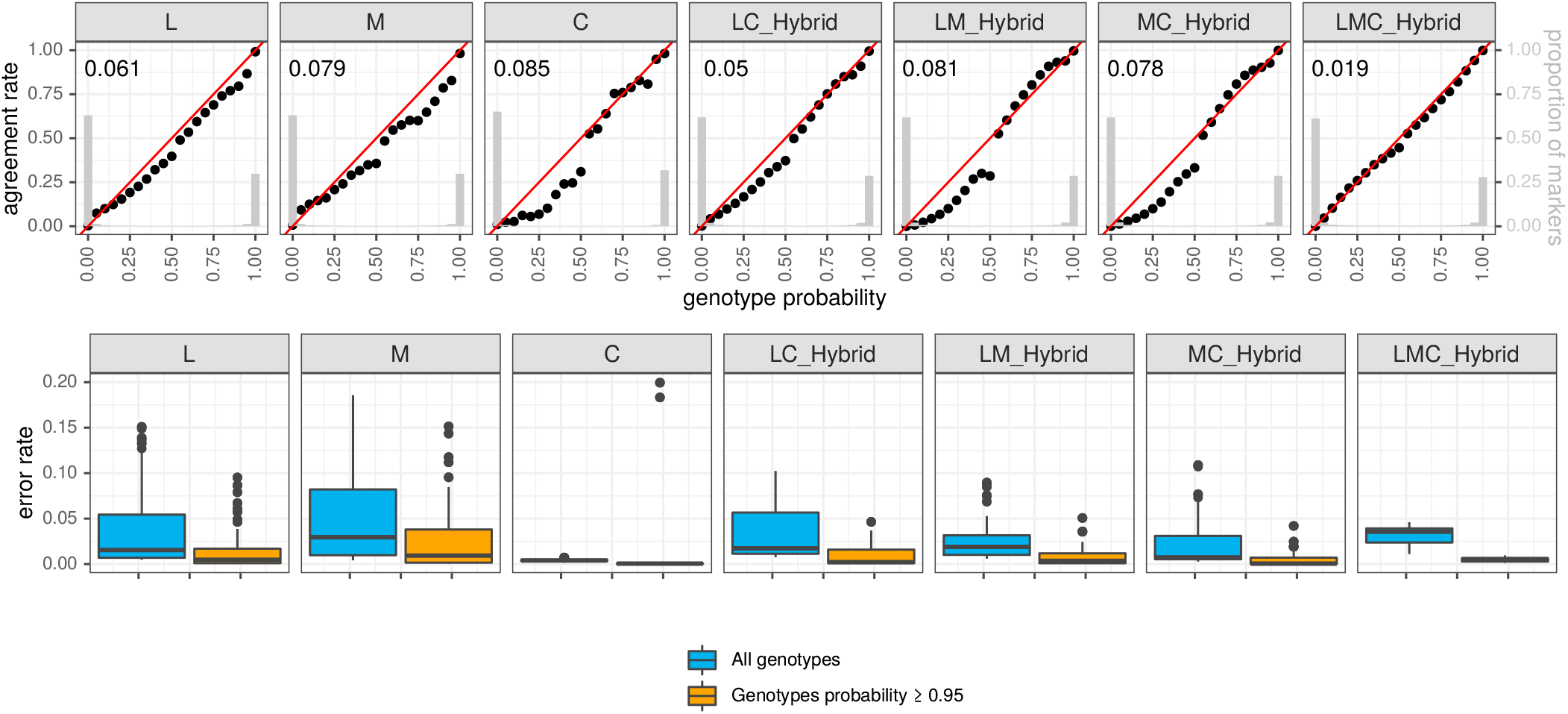
Queen genotype reconstruction for each of the seven groups. Detailed genotype calibration and genotyping error rate for each group. The first row represents the genotype calibration, with AUC and genotype probability distribution, for each of the groups tested when performing queen genotype reconstruction using simulations from real data on the whole genome are clustering on genetic ancestries estimated with AM. The second row represents the genotyping error rate for each of the scenarios tested when performing queen genotype reconstruction using simulations from real data on the whole genome for the whole genome or after filtering on best genotype probability equal to or greater than 0.95.

**Figure S6:**
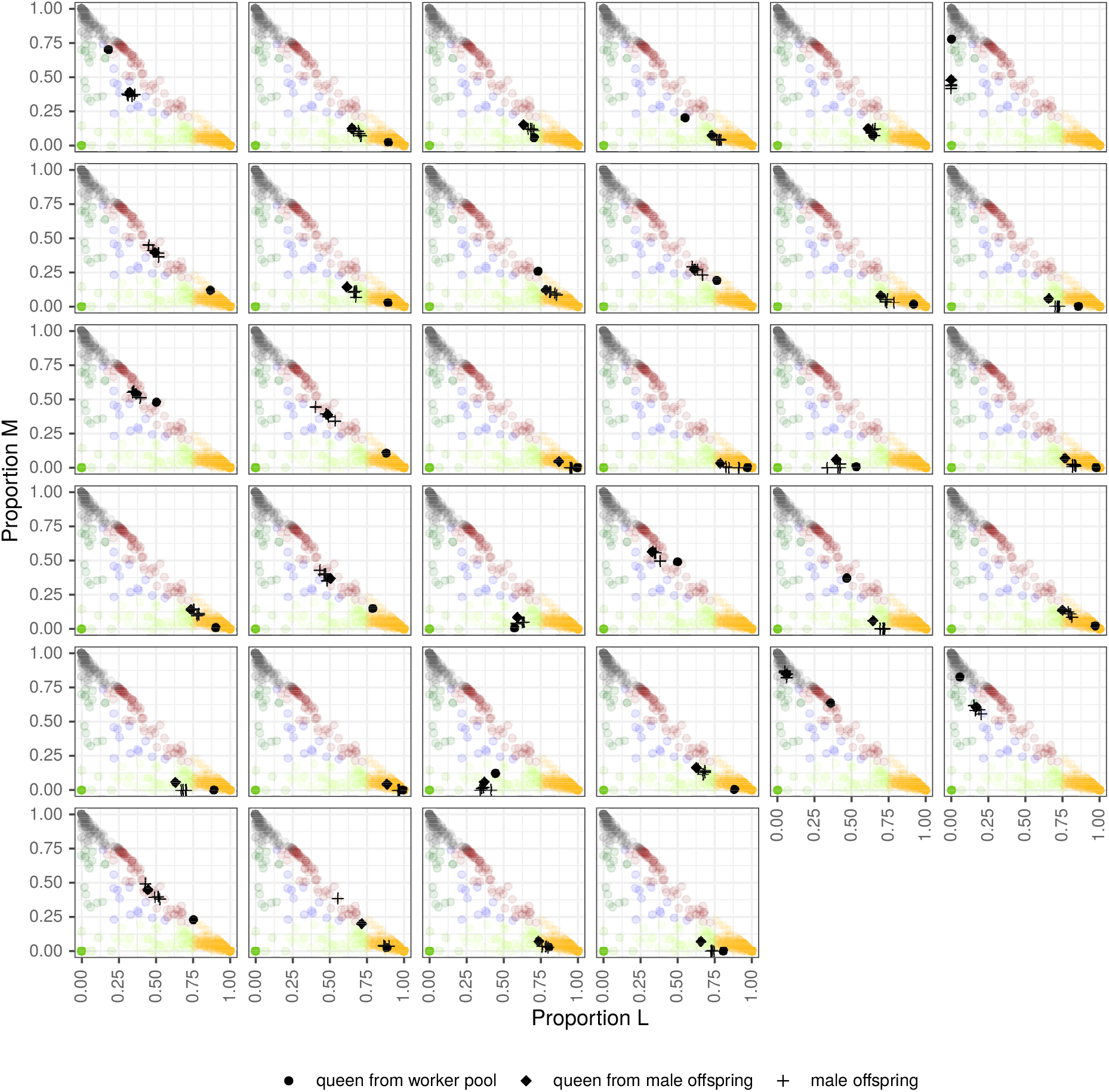
Genetic ancestries on experimental colonies estimated from different models and data on experimental colonies. Two dimensions plot of genetic ancestries for the different estimates on the experimental colonies. For the 34 experimental colonies, drones offspring of the queen (crosses), queen reconstructed from these drones (diamonds) and queen reconstructed from the pool experiment (circles) projected on top of the individuals from the diversity panel (628 from Wragg et al. (2021)), representing genetic ancestries in two dimensions.

**Figure S7:**
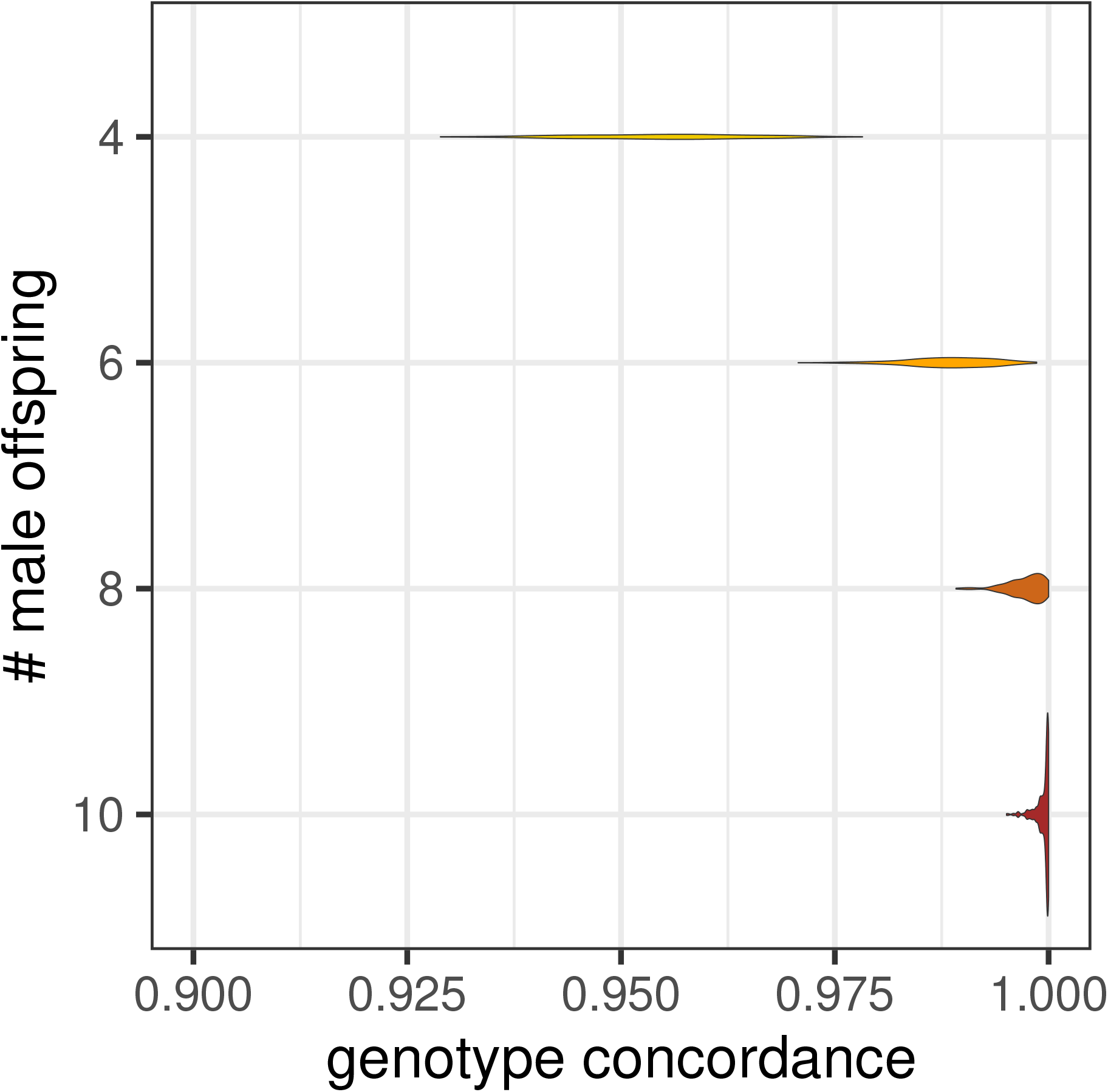
textbfConcordance between real and reconstructed queen genotypes as a function of the number of male offspring available

**Figure S8:**
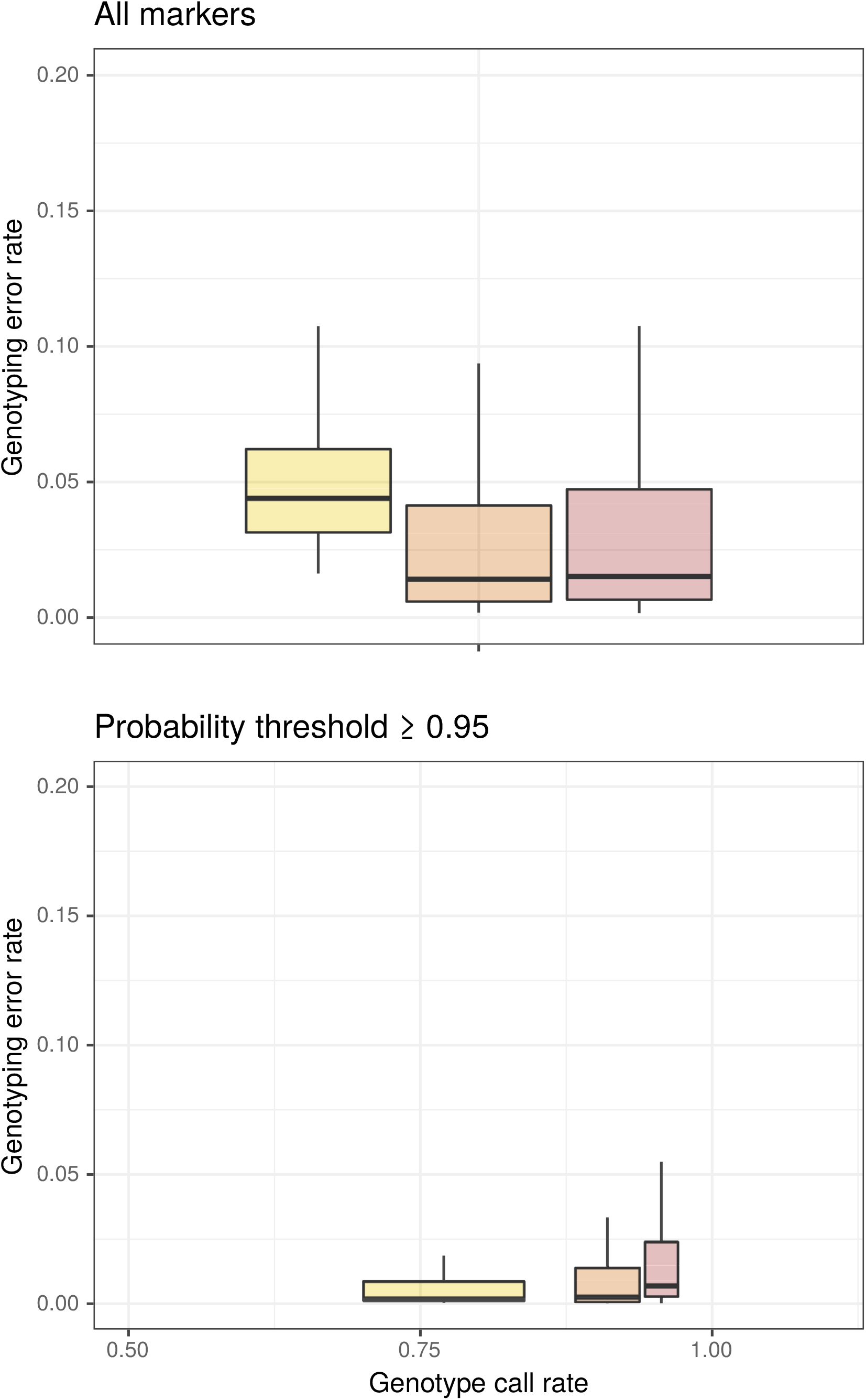
Genotyping error rates for different sequencing depths. Impact of pool sequencing depth on genotyping error rate. Genotyping error across each colony simulated for linked markers across the whole genome after genotype reconstruction within groups of homogeneous genetic ancestries based on estimations from AM for depth 10 (yellow), 30 (orange) and 100 (brown). The top panel is for all markers on the genome, the bottom panel is for markers with best genotype probability higher or equal than 0.95, the x axis represents the genotype call rate.

## References

Alexander, D. H., Novembre, J., & Lange, K. (2009). Fast model-based estimation of ancestry in unrelated individuals. Genome Research, 19, 1655–1664. doi:10.1101/gr.094052.109

Bansal, V. (2010). A statistical method for the detection of variants from next-generation resequencing of dna pools. Bioinformatics (Oxford, England), 26(12), i318–i324. doi:10.1093/bioinformatics/btq214

Brascamp, E. W. & Bijma, P. (2014). Methods to estimate breeding values in honey bees. Genetics Selection Evolution, 46(1), 53. doi:10.1186/s12711-014-0053-9

Chang, C. C., Chow, C. C., Tellier, L. C. A. M., Vattikuti, S., Purcell, S. M., & Lee, J. J. (2015). Second-generation plink: Rising to the challenge of larger and richer datasets. GigaScience, 4(1). doi:10.1186/s13742-015-0047-8

Gautier, M., Vitalis, R., Flori, L., & Estoup, A. (2021). F-statistics estimation and admixture graph construction with pool-seq or allele count data using the r package poolfstat. submitted. doi:10.1101/2021.05.28.445945

Hivert, V., Leblois, R., Petit, E., Gautier, M., & Vitalis, R. (2018). Measuring genetic differentiation from pool-seq data. Genetics, 210(1), 315–330. doi:10.1534/genetics.118.300900. eprint: https://www.genetics.org/content/210/1/315.full.pdf

Jones, J. C., Du, Z. G., Bernstein, R., Meyer, M., Hoppe, A., Schilling, E., … Bienefeld, K. (2020). Tool for genomic selection and breeding to evolutionary adaptation: Development of a 100k single nucleotide polymorphism array for the honey bee. Ecology and Evolution, 10(13), 6246–6256. doi:https://doi.org/10.1002/ece3.6357. eprint: https://onlinelibrary.wiley.com/doi/pdf/10.1002/ece3.6357

Kofler, R., Pandey, R. V., & Schlötterer, C. (2011). Popoolation2: Identifying differentiation between populations using sequencing of pooled dna samples (pool-seq). Bioinformatics, 27(24), 3435–3436. doi:10.1093/bioinformatics/btr589

Li, H. (2013). Aligning sequence reads, clone sequences and assembly contigs with bwa-mem. arXiv preprint arXiv:1303.3997.

Li, H. & Durbin, R. (2009). Fast and accurate short read alignment with burrows–wheeler transform. Bioinformatics, 25(14), 1754–1760. doi:10.1093/bioinformatics/btp324

Liu, H., Zhang, X., Huang, J., Chen, J. Q., Tian, D., Hurst, L. D., & Yang, S. (2015). Causes and consequences of crossing-over evidenced via a high-resolution recombinational landscape of the honey bee. 16(1), 15. Retrieved from https://doi.org/10.1186/s13059-014-0566-0

Micheletti, S. J. & Narum, S. R. (2018). Utility of pooled sequencing for association mapping in nonmodel organisms. Molecular Ecology Resources, 18(4), 825–837. doi:https://doi.org/10.1111/1755-0998.12784

Oxley, P. R. & Oldroyd, B. P. (2010). The genetic architecture of honeybee breeding. 39, 83–118.

Petersen, G. E. L., Fennessy, P. F., Van Stijn, T. C., Clarke, S. M., Dodds, K. G., & Dearden, P. K. (2020). Genotyping-by-sequencing of pooled drone dna for the management of living honeybee (apis mellifera) queens in commercial beekeeping operations in new zealand. Apidologie. doi:10.1007/s13592-020-00741-w

Pritchard, J. K., Stephens, M., & Donnelly, P. (2000). Inference of population structure using multilocus genotype data. Genetics, 155(2), 945–959. Retrieved from %3CGo%20to%20ISI%3E://WOS:000087475100039

Purcell, S., Neale, B., Todd-Brown, K., Thomas, L., Ferreira, M. A. R., Bender, D., … Sham, P. C. (2007). Plink: A tool set for whole-genome association and population-based linkage analyses. American journal of human genetics, 81(3), 559–575. doi:10.1086/519795

Saelao, P., Simone-Finstrom, M., Avalos, A., Bilodeau, L., Danka, R., de Guzman, L., … Tokarz, P. (2020). Genome-wide patterns of differentiation within and among u.s. commercial honey bee stocks. BMC Genomics, 21(1), 704. doi:10.1186/s12864-020-07111-x

Schlötterer, C., Tobler, R., Kofler, R., & Nolte, V. (2014). Sequencing pools of individuals — mining genome-wide polymorphism data without big funding. 15, 749. Retrieved from http://10.1038/nrg3803

Speed, D., Holmes, J., & Balding, D. J. (2020). Evaluating and improving heritability models using summary statistics. Nature Genetics, 52(4), 458–462. doi:10.1038/s41588-020-0600-y

Tarpy, D. R. & Nielsen, D. I. (2002). Sampling error, effective paternity, and estimating the genetic structure of honey bee colonies (hymenoptera: Apidae). Annals of the Entomological Society of America, 95(4), 513–528. doi:10.1603/0013-8746(2002)095[0513:SEEPAE]2.0.CO;2

Tarpy, D. R., Nielsen, R., & Nielsen, D. I. (2004). A scientific note on the revised estimates of effective paternity frequency in apis. Insectes Sociaux, 51(2), 203–204. doi:10.1007/s00040-004-0734-4

Toth, A. L. & Zayed, A. (2021). The honey bee genome–what has it been good for? Apidologie. doi:10.1007/s13592-020-00829-3

Uzunov, A., Brascamp, E. W., & Büchler, R. (2017). The basic concept of honey bee breeding programs. Bee World, 94(3), 84–87. doi:10.1080/0005772X.2017.1345427

Wallberg, A., Schöning, C., Webster, M. T., & Hasselmann, M. (2017). Two extended haplotype blocks are associated with adaptation to high altitude habitats in east african honey bees. PLOS Genetics, 13(5), e1006792. doi:10.1371/journal.pgen.1006792

Wallberg, A., Bunikis, I., Pettersson, O. V., Mosbech, M. B., Childers, A. K., Evans, J. D., … Webster, M. T. (2019). A hybrid de novo genome assembly of the honeybee, apis mellifera, with chromosome-length scaffolds. BMC Genomics, 20(1), 275. doi:10.1186/s12864-019-5642-0

Wragg, D., Marti-Marimon, M., Basso, B., Bidanel, J. P., Labarthe, E., Bouchez, O., … Vignal, A. (2016). Whole-genome resequencing of honeybee drones to detect genomic selection in a population managed for royal jelly. 6, 27168. Retrieved from https://www.nature.com/articles/srep27168#supplementary-information

Wragg, D., Eynard, S. E., Basso, B., Canale-Tabet, K., Labarthe, E., Bouchez, O., … Vignal, A. (2021). Complex population structure and haplotype patterns in western europe honey bee from sequencing a large panel of haploid drones. bioRxiv, 2021.09.20.460798. doi:10.1101/2021.09.20.460798

Zhou, X. & Stephens, M. (2012). Genome-wide efficient mixed-model analysis for association studies. Nature Genetics, 44(7), 821–824. doi:10.1038/ng.2310

